# Identification of key bacteria for ecological dynamics in a coastal marine observatory

**DOI:** 10.1101/2025.06.16.659933

**Authors:** Erick Mateus-Barros, Emiliano Pereira, Juan Zanetti, Luciana Griffero, Rudolf Amann, Cecilia Alonso

**Affiliations:** Grupo de Ecología Microbiana de Sistemas Acuáticos, Centro Universitario Regional del Este, Universidad de la República. Uruguay; Molecular Ecology Department, Max Planck Institute for Marine Microbiology, Bremen Germany

**Keywords:** marine bacterioplankton, temporal dynamics, amplicon, CAZimes, biological interactions

## Abstract

Bacteria are essential for ecosystem maintenance, drive biogeochemical cycles, and influence responses to climate change. Additionally, many are involved in blooms and toxin production, affecting environmental quality. Despite their crucial role, some of their characteristics are only now being understood with the development of molecular techniques independent of cultivation. Thus, the recent effort to unify classical and microbial ecology studies has led to a better understanding of their dynamics. Here, we investigate the temporal processes that influence bacterial abundance and persistence, as well as their role in promoting or inhibiting the presence of certain organisms in the core of a metacommunity. Using molecular approaches such as amplicon and metagenomic analyses, we retrieved information on key bacteria (those with high abundance and/or persistence) from the South Atlantic Microbial Observatory (SAMO) to assess their interactions with environmental and biological factors. Bimodality was hardly observable, while the community structuring reflected the influence of large scale environmental processes in a recurrent pattern that clusters winter-spring and summer-autumn communities, reinforcing previous findings of intense influence of marine currents in this region. Among the key organisms, SAR11 and Flavobacteriaceae stand out. Flavobacteriaceae is known for its adaptive capacity and large genomes, while SAR11 has a reduced genome, making it dependent on compounds produced by other organisms. We identified an annual fluctuation in its abundance linked to Synechococcus blooms. The significant co-occurrence of ASVs from both groups reinforces the evidence of a biological interaction that sustains SAR11 through the exchange of these compounds. Through this study, we contribute to clarifying the factors that locally influence the geographic patterns of marine bacteria and identifying the pathways that promote their dynamics and functions in the ecosystem.

## Introduction

Bacteria are essential for diverse ecological functions and the structure of communities. Being the most abundant organisms in any ecosystem they are key for ecosystems maintenance (Chiriac et al. 2022), driving the major biogeochemical cycles (Myklestad 2000), and influencing the responses of ecosystems to climate change (Cavicchioli et al. 2019). In addition, many of them are related to phenomena such as blooms and toxin production (Huisman and Hulot 2005), which affect environmental quality. Despite their crucial environmental roles, some of their characteristics could be fully understood only after the advent of culture-independent molecular techniques (Rappé and Giovannoni 2003), and key aspects of their ecology remain still to be addressed. For example, recent studies on microbial biogeography have shown clearly identifiable patterns of dispersion and isolation for these organisms (Martiny et al. 2006, Lindh et al. 2017, Soininen et al. 2018, Mateus-Barros et al. 2021), which is contrary to what was initially supposed based on their small size and large populations.

In this context, there is a running effort to unify studies in classical and microbial ecology (Horner-Devine and Bohannan 2006) to boost the development of both disciplines (Acinas et al. 2004, Barberán et al. 2014). On one hand, this interaction allows us to apply mathematical models and ecological theories to microbial ecology (e.g. Lindh et al. 2017, Soininen et al. 2018); on the other hand, microbial datasets represent an ideal sample number to test the limits of ecological theories (Barberán et al. 2014).

The core-satellite hypothesis (Hanski 1982) is a conceptual framework of classical ecology that emerges as an attractive way to explore the macroecological structure of microorganisms (Lindh et al. 2017, Jeong et al. 2020, Mateus-Barros et al. 2021, Escalas et al. 2022). This hypothesis states that the frequency of occupancy in a metapopulation should be bimodal. This spatial pattern of species distribution indicates that, when the sampling effort is ideal, most organisms found should be present in one or few sites, while others will be distributed throughout the entire landscape (Gleason 1929). This distribution dichotomy evidences the existence of two distinct groups of organisms regarding occupancy: on the one hand, there are the satellites which are those restricted to one or few sites, while on the other, the most persistent organisms are called core (Hanski 1982). This phenomenon seems to be mainly related to abundance variation through space and time since these organisms are generally those that manage to (re)colonize new sites in a dispersal wave that gradually increases the area covered by a species (Brown 1984, Papp and Izsák 1997).

Utilizing the core-satellite framework it is possible to recover some interesting information about biodiversity since its pattern depends on the dispersion capacity inherent to each organism that should impact this structural feature. For example, some aquatic bacteria can be regionally persistent even when they are not abundant (Lindh et al. 2017). Additionally, while the satellite encompasses the major part of regional diversity (Mateus-Barros et al. 2021), the core is composed of organisms that are potentially key to ecosystem functions (Fonte et al. 2021). The core-satellite approach is also a valuable way to retrieve information concerning the metacommunity structure (i.e. a set of local communities linked by a flux of individuals). Also, this approach can recover details on topics such as colonization and extinction dynamics, adaptiveness and dispersal capacity, niche partition, and competition (Gaston et al. 2000, Mehranvar and Jackson 2001, Lindh et al. 2017). Despite its broad applicability and ease of implementation, this approach remains rarely employed in marine microbial ecology studies and has only rarely been applied in environments located in the Northern Hemisphere (Lindh et al. 2017).

Therefore, applying these approaches to large-scale temporal data can provide new insights into the dynamics of marine microbial communities and their interactions with environmental factors and other biotic components. Locally, environmental factors like pH, temperature and salinity (Lindström et al. 2005, Sunagawa et al. 2015) and large scale processes like shifts in marine currents domains or seasonal conditions (Messer et al. 2020, Pereira et al. 2025) are expected to determine which groups will likely be more abundant and, consequently, have increased chances to disperse a highest number of individuals and dominate spatially (Gaston et al. 2000), having consequences at the ecosystem scale. For example, a long-term study on the dynamics of a phytoplanktonic community in a tropical lake showed that the change in environmental conditions changed dominance features, and allowed a non-seasonal change of ecological state, from filamentous cyanobacteria to green algae (Rugema et al. 2019). Biological interactions can also be a relevant feature to be considered. A microbial community is connected in an intricate interaction web (i.e. microbial loop Azam et al. 1983) by compounds incorporation made available by exudation, excretion, and cell lysis by other organisms (Kawasaki and Benner 2006, Alonso et al. 2012, Sarmento and Gasol 2012). Despite this, it remains unclear which components of the microbial community are most strongly affected by these variables. Complementary, there is a lack of information on the temporal dynamics of marine microbial communities in the Southern Hemisphere (Fermani et al., 2024), which hinders more informed decision-making in various areas, including conservation and management.

In this sense, we sought to investigate microbial temporal dynamics in the South Atlantic Microbial Observatory (SAMO), a microbial observatory located at the uruguayan coast, in a region characterized by fluctuations of distinct physico-chemical factors operating at different temporal scales, with the potential to strongly impact the dynamics that structure the community. For example, In a recent study conducted in this observatory (Pereira et al., 2025), it was demonstrated that the temporal structuring of this microbial community is strongly structured by successive seasonal variation guided by the confluence of Brazil and Malvinas currents (Burone et al. 2021). It includes a change of composition that turns this community more similar to those found in sites uniquely influenced by one of each current (Pereira et al., 2025). Thus, this Microbial Observatory emerges as an exceptional site for observing how these distinct processes may be interconnected, influencing local dynamics of persistence and dominance in marine microbial communities of the Southern Hemisphere.

More specifically, our aim was to highlight the local temporal processes that may contribute to the abundance and persistence of bacteria, and their roles in guaranteeing or preventing certain organisms from being included in the core of a metacommunity. More specifically, we intend to (1) assess how temporal variation on the bacterial presence and abundance correlates with environmental conditions and the dynamics of other biological components, and to (2) identify the potential roles of the more abundant taxa by analyzing their associations with environmental variables, other organisms, and genes related to the carbon cycle.

## Methods

### Study Sites and Sampling Design

In this study we used a dataset collected monthly in the South Atlantic Microbial Observatory (SAMO) located in the marine zone of the ―Laguna de Rocha‖ protected area (34° 42‘ S, 54° 15‘ W) to test the impact of a series of local environmental and biological variables on the persistence and abundance variation of bacteria aiming to understand what makes them have a core or satellite distribution along time.

The SAMO, launched in 2014 from the Ocean Sampling Day international initiative (Kopf et al. 2015), is located at the Uruguay–Buenos Aires shelf ecoregion. This marine region is categorized by having a complex hydrodynamic system, as it is impacted by the Río de la Plata outflow and the confluence of Brazil and Malvinas currents (Burone et al. 2021). This creates an intricate balance of temperature, salinity, and nutrients that varies through the year depending on the water mass that is predominantly influencing the region (Burone et al. 2021). While the warmer and salty Brazil current mainly impacts the region duringthe Winter-Spring period, the Malvinas current turns the ocean colder in the summer-Autumn period (Burone et al. 2021).

The SAMO is part of the Latin American Aquatic Microbial Observatory Network (AMOLat), a scientific consortium for the creation and maintenance of long-term microbial observatories all around the continent (Fermani et al. 2024).

### Sampling procedures, physicochemical and molecular analyses

The sampling of sub-surface marine water has been performed monthly since 2018 to nowadays. In the field, a multi-parameter probe is used to recover environmental information (salinity, temperature, pH, turbidity, dissolved oxygen, suspended solids, and fluorescence). Further physicochemical characterization includes: i) the quantification of nutrients (total phosphorus, total nitrogen, phosphate, ammonium, nitrite, nitrate, reactive silica) and chlorophyll a, following standard methods (Apha 1999), and ii) the spectral characterization of the chromophoric and fluorescent dissolved organic matter (cDOM and fDOM) (Alonso et al under revision).

For the molecular analyses, departing from the microbial biomass recovered after water samples filtration, the DNA was extracted following standard methods (Alonso et al 2010). The prokaryotic community diversity was determined by amplifying a variable region of the 16S rRNA gene with the primers 515F-Y (5‘-GTG YCA GCM GCC GCG GTA A-3‘) and 916R (5‘-CCG YCA ATT YM TTT RAG TTT-3‘) (Parada et al. 2015), while the small planktonic eukaryotic components were determined by amplifying a variable region of the 18S rRNA gene using the primers TAReukFWD1 (5’-CCAGCASCYGCGGTAATTCC-3’) and TAReukREV3 (ACTTTCGTTCTTGATYRA) (Stoeck et al. 2010).

The 16S and 18S amplified material was sequenced on an Illumina MiSeq at the Integrative Genomics Core (City of Hope, US). The extracted DNA without prior amplification was sequenced on the Illumina NovaSeq 6000 SP FC platform at LGC Genomics GmbH (Berlin, Germany) and the Genomics and Cell Characterization Core Facility (GC3F) at the University of Oregon (Eugene, US) to obtain the prokaryotic, eukaryotic, and metagenomic sequences of the bacterioplankton community, respectively.

### Bioinformatics

The amplicon sequence data were processed with the DADA2 pipeline (Callahan et al. 2016), integrating the following tasks: removing primers, trimming low-quality sequence regions, dereplicating, identifying amplicon sequence variants (ASVs), merging paired-end reads, and removing chimeras. Prior to the analyses, the ASVs tables were cleaned to remove sequences non-identified as bacteria and eukaryotes in the 16S and 18S datasets, respectively. Afterwards, the tables were rarefied by the sample with lowest sequence number, and the ASVs counting five or fewer sequences were removed.

To pre-process the metagenomic data, we applied the following approach: remotion of adapter sequences, merge of paired-end reads, quality trimming and filtering of merged and unmerged reads. The preprocessed metagenomic data was analyzed utilizing a custom-developed pipeline (*i.e.,* Mg-Traits, Pereira-Flores et al. 2021), which performs the computation of several functional aggregated traits at the metagenome level, ranging from GC variance to average genome size, and includes the functional annotation based on the Pfam database (Mistry et al. 2021) and the carbohydrates-active enzymes database (Drula et al. 2022), among others. The results of the metagenomic sequence data analysis are represented as Functional Abundance Matrices (FAMs). In addition, a matrix summarizing the metagenomic trait values computed for each metagenomic sample is generated. From all this data that was generated, we had filtered the contigs that decode the proteins related to the Carbon cycle (here after they will be called CAZymes).

In order to ensure better representativeness of samples collected across different seasons over the years, we performed a subsampling by selecting 35 of the samples collected between 2018 and 2021.

### Statistical analyses

As stated before, the occupancy capacity of an organism seems to be related to their abundance features in each location (Papp and Izsák 1997), as those more abundant tend to more easily (re)colonize other sites in a landscape (Brown 1984). That being said, bacteria have a remarkably high dispersal capacity in nature, which means that some groups can reach longer distances while maintaining a low mean relative abundance (Lindh et al. 2017). In this sense, to better understand the bacterioplanktonic temporal dynamics observed in SAMO, we carried out statistical measurements that consider the entire community and sub-sampled the community into those ASVs considered as key, by observing two main characteristics, *i.e.* mean relative abundance and persistency. Each ASV‘ abundance was classified from the approach proposed by Pedrós-Alió (2012), which defined as abundant those organisms which reach at least 1% of local relative abundance. Persistence was defined as the capacity to be present in more than 90% of all samples, in an approach similar to that proposed by Hanski (1982) to classify organisms that better occupy a landscape. Considering this, we defined 3 main categories: ASVs with a mean relative abundance greater than 1% but present in less than 90% samples were classified as ―Abundant‖; conversely ASVs with mean relative abundance below 1%, but present in at least 90% of samples persistence were called ―Persistent‖. The ―Core‖ of bacterioplankton comprised ASVs with both high abudandance (>1%) and high persistence (>90%). Finally, ASVs that did not meet either the Abundance or Persistence thresholds were classified as ―non-Key‖.

To determine whether there was a bimodal distribution on the persistence of microorganisms, we applied the Mitchell-Olds & Shaw test of quadratic extremes (Mitchell-Olds and Shaw 1987). A significant bimodal distribution indicates the presence of a group of ASVs always present in SAMO independently of the environmental conditions, while other groups appear only under specific circumstances (Gleason 1929, Hanski 1982).

To determine the relationship between the temporal variations of bacteria abundances and environmental variables, and corroborate this pattern in a wider time-span, we applied the non-Metric Multidimensional Scaling (nMDS) and distance-based Redundancy Analysis (dbRDA). To understand how the community dissimilarity fluctuated through the years, we applied a Time-decay Relationship (TdR) analysis. Complementerly, to assess potential direct and indirect biological relationships between the prokaryotes classified as key in this community and both with eukaryotes and key functions related to C cycle, we used paired co-occurrence analysis. For this purpose, we combined the 16S and 18S data with the metagenomic contigs containing information on the CAZymes family synthesis. All statistical analyses described above were performed using the R software (R Core Team 2022) and various functions provided by the vegan (Oksanen et al. 2016) and Tidyverse (Wickham et al. 2019) packages. The co-occurrence analysis was performed with the co-occur package (Griffith et al. 2016).

## Results

After rarefaction and removal of the rarest ASVs, 2,586 ASVs were retained from a total of 326‘540 reads (mean of 9,330 reads per sample), from which we identified 30 phyla. These ASVs were classified — when possible at the Order, Family, or Genus level — into 31 groups of interest. The most abundant groups of interest encompassed the SAR11 (17.75% of total reads), followed by Flavobacteriaceae (14.77%), Rhodobacterales (8.79%), other Gammaproteobacteria (6.05%), Cryomorphaceae (5.85%), and Synechococcus (5.32%) (Fig. 1).

**Figure 1.**
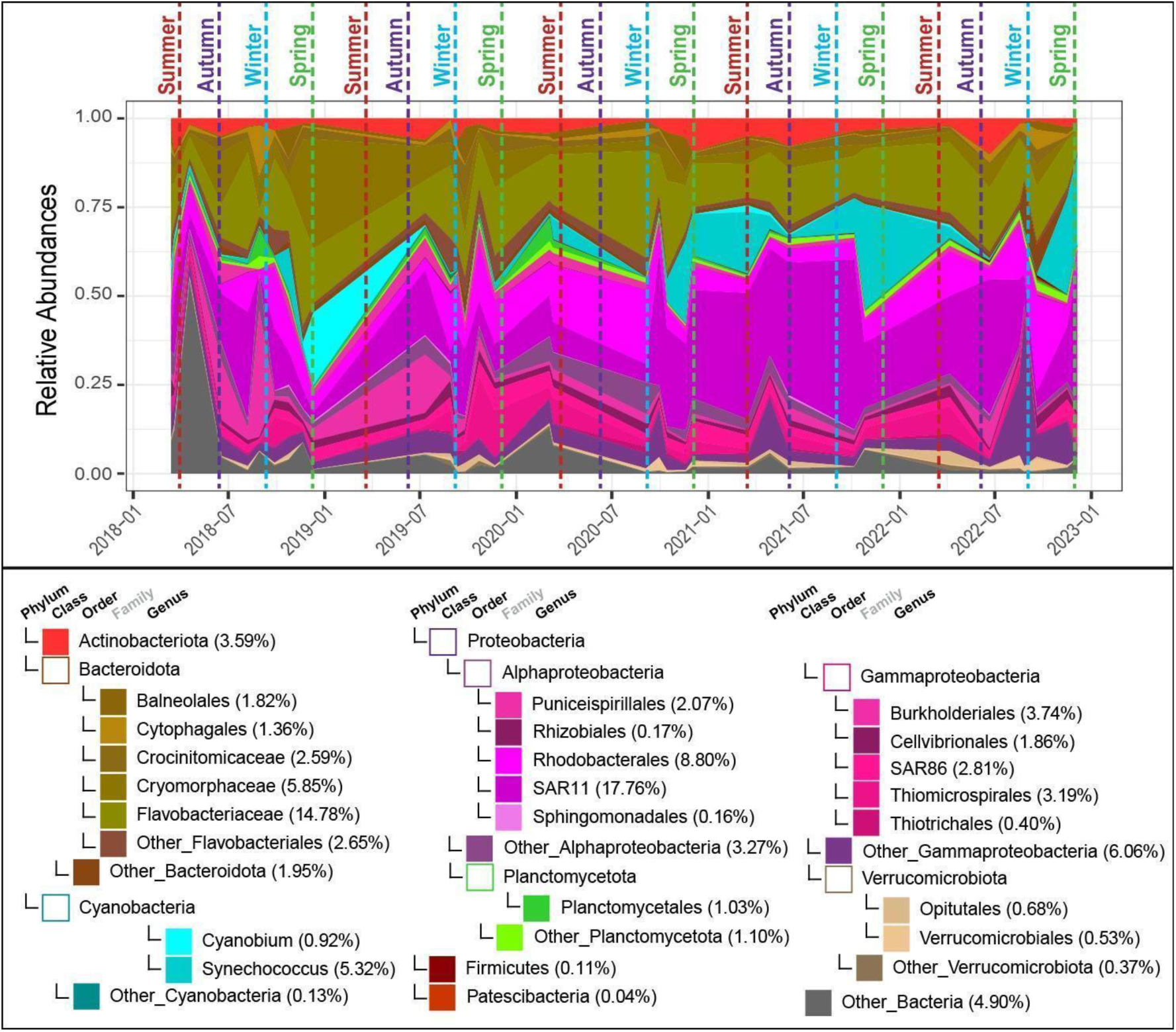
Temporal variation in the relative abundance of bacterioplankton. Each ASV was assigned to a phylogenetic phylum. Groups with known relevance to local dynamics were classified at the Class, Order, or Genus level, depending on the level of taxonomic resolution achieved for each phylum. Parenthetical values represent the total relative abundance of each group

Despite the bimodality test returned significant results (p > 0.05), it is hardly seen (Fig. S1). To better understand what may be driving this phenomenon, we decomposed the data into a heatmap showing the fluctuations in abundance for each ASV identified as key (*i.e.*, Abundant, Persistent, and Core) over the years (Fig. S2). Here, we can observe variation in the abundance and persistence of these ASVs, which prevents more consistent persistence of these taxa. In any case, this variation does not exhibit a high Bray-Curtis dissimilarity.

Additionally, we observed a significant correlation between community variation and specific environmental and biological components. The nMDS analysis (Fig. 2) shows that large-scale variations in the community appear to be related to seasonal patterns that divide the samples into two main groups by season: Summer-Autumn and Winter-Spring, in a recurrent arrangement that does not vary considerably with interannual changes, but is also reflected in the observed diversity variations between seasons. We also found a trophic state effect on this observed temporal variation.

**Figure 2.**
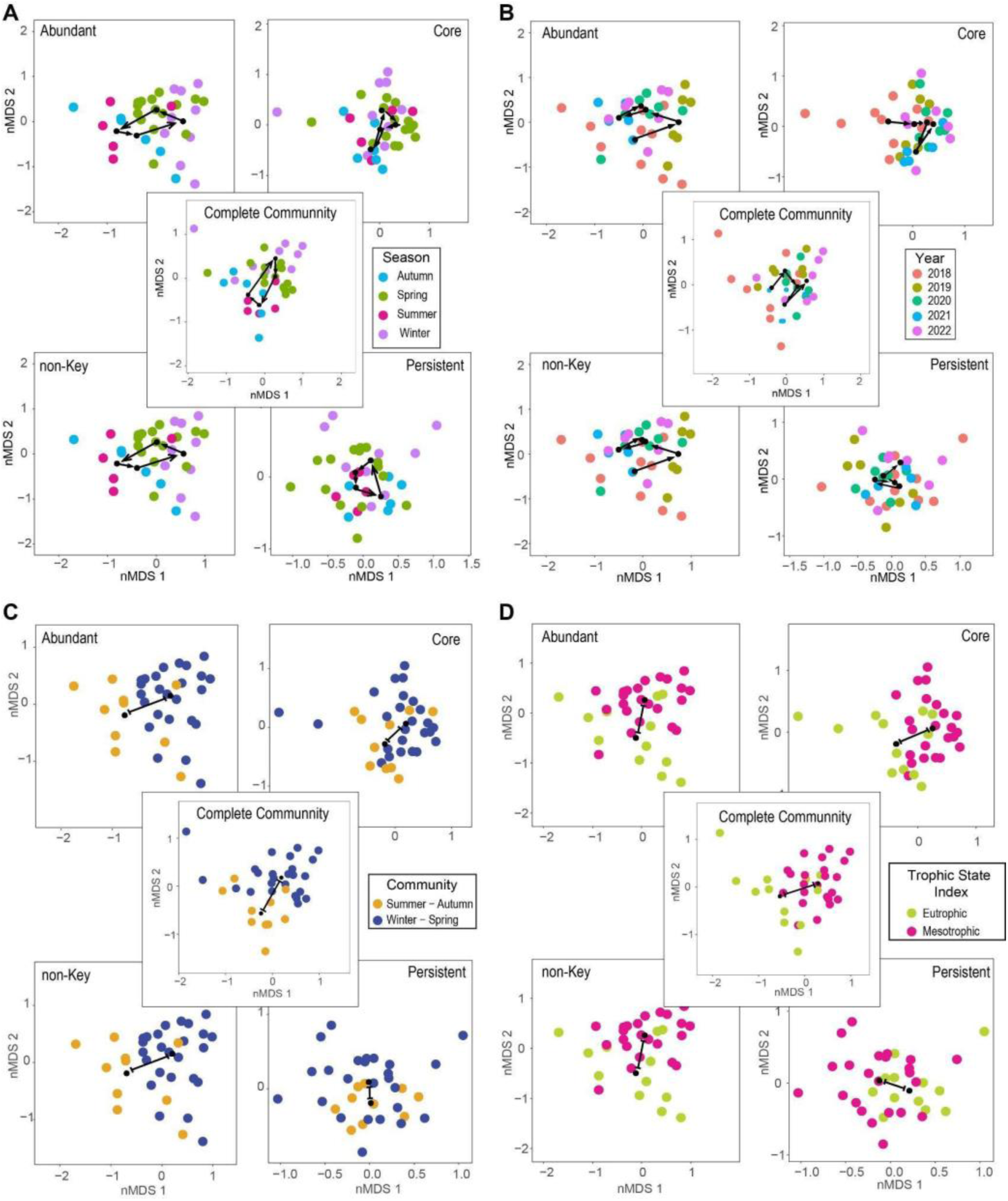
nMDS (non-Metric Multidimensional Scaling) plots depicting the bacterioplankton community dissimilarity (stress = 0.1513) and decomposed into different key groups – Core (stress = 0.1626), Abundant (stress = 0.1916), Persistent (stress = 0.1846), and non-key (stress = 0.1382) for each of the large-scale environmental changes taken into account. Seasonal changes (A), year of collection (B), the winter-spring to summer-autumn transition by the Brazil-Malvinas Confluence (C), and trophic state (D) were considered

The beta-dispersion analysis (Fig. S4) indicated that community variation was not significantly related to within-group dispersion. Further insights were gained by subdividing the community into key bacteria (Core, Abundant, and Persistent) and non-key bacteria. In the Abundant and non-Key ASV groups, we observed a degree of variation comparable to that of the full community. In contrast, Core and Persistent ASVs displayed lower dispersion and more compact clustering patterns (Fig. 2), consistent with their expected ecological stability.

Complementarily, the dbRDA (Fig. 3A) illustrates that temporal variation in microbial community composition is significantly associated with environmental gradients (p = 0.007). The first axis (explaining 28.75% of the variation) is strongly associated with temperature, indicating its dominant role in seasonal community shifts. The second axis (19.80% of variation) reflects gradients primarily related to salinity, nitrogen compounds (DIN and Total Nitrogen), and protein-like substances (Component 4). Additionally, this axis captures variation related to the Trophic State Indices for phosphorus (TSI–P), chlorophyll-a (TSI–Chla), and water transparency (TSI–Secchi). Dissolved Oxygen shows an inverse relationship with many of these environmental drivers. Subsampling the key and non-key groups revealed that, while salinity and nutrient-related variables remained the main drivers of community variation, each group exhibited distinct associations with environmental gradients (Fig. 3B). In the non-Key group (p = 0.003), salinity and nutrient concentrations (DIN, TN) were the predominant drivers, with pH and temperature also contributing as secondary factors. In the Abundant group (p = 0.174), variation was mainly aligned with two contrasting gradients: one associated with salinity and temperature, and the other with nutrient availability and productivity proxies. Persistent ASVs (p = 0.808) showed more diffuse structuring, suggesting three subclusters with distinct associations: one linked to salinity, another to dissolved oxygen, and a third to nutrients and turbidity. The Core group (p = 0.003) exhibited a strong and simultaneous relationship with salinity, temperature, and nutrient variables, including FDOM Components 3 and 5 — interpreted as protein-like fractions — suggesting a closer coupling with biologically labile organic matter. Component 4 — described as terrestrial humic substances — was consistently associated with all groups, suggesting a broad influence of allochthonous organic matter on microbial community structure across community components.

**Figure 3.**
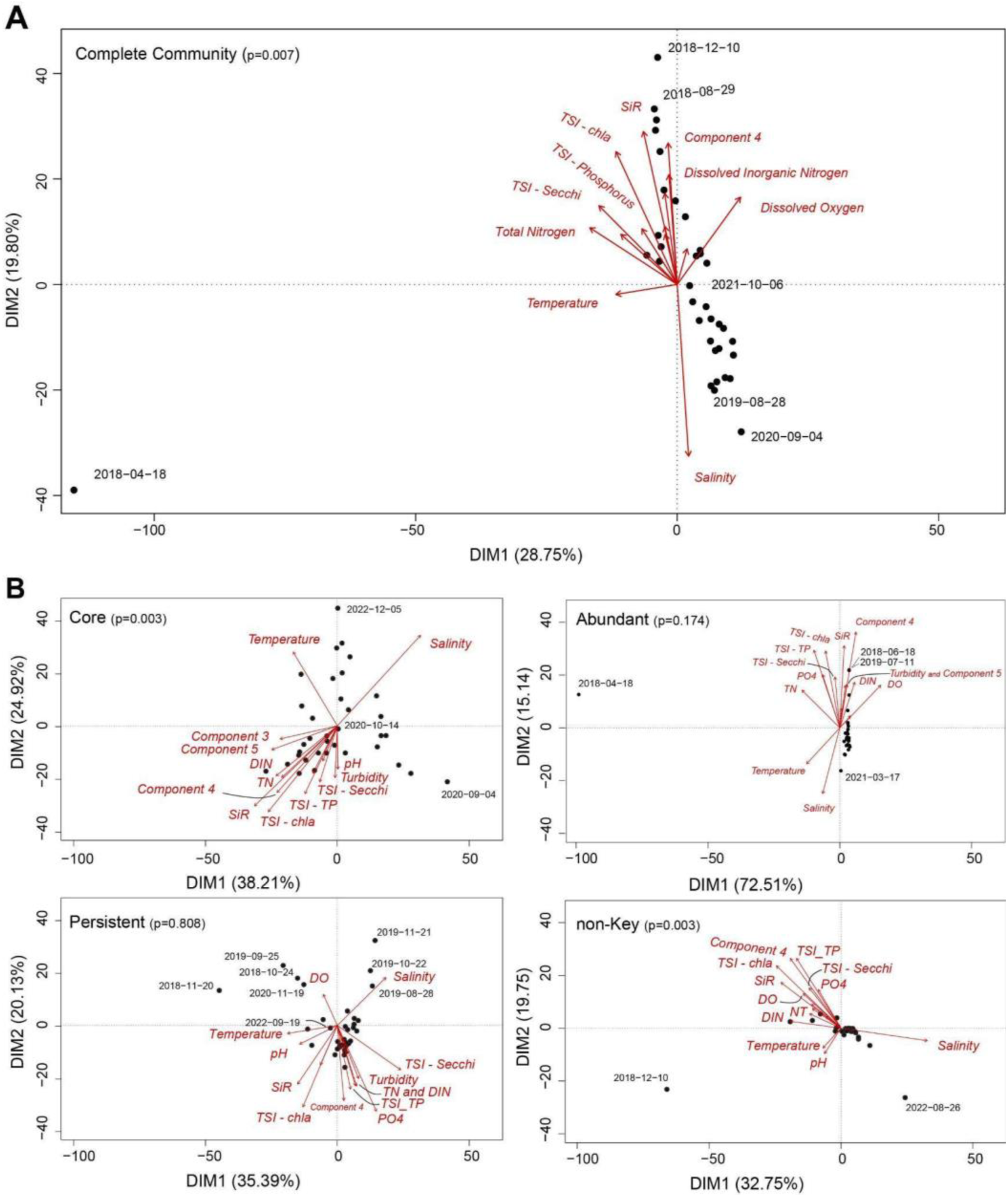
dbRDA (distance-based ReDundancy Analysis) plots displaying the maximum similarity among sites, considering the relationship between them and the environmental factors that best explain the observed bacterioplankton dissimilarity. The complete community (A) was decomposed into their key groups (B): Core (upper left box), Abundant (upper right box), and Persistent (lower left box), and non-key (lower right box) of the observed bacterioplankton. The results were significant (p < 0.05) for the ASVs of the Complete Community, as well as the Core and non-key groups

The TdR analysis shows how community dissimilarity increases with the temporal distance between samples (Fig. 4). As expected, a gradual increase in dissimilarity is observed, displaying a characteristic wave-like pattern with a one-year periodicity. This indicates that samples collected in the same season across different years tend to be more similar, likely reflecting the recurrence of seasonal environmental conditions—a pattern consistent with what was observed in Fig. 2. . This trend, observed in the community as a whole, is more prominent in the subset of Abundant and non-Key groups, while the variation is less evident among Core bacteria and absent in the Persistent ones. These two groups exhibit lower temporal variability and minimal rates of change even over longer time intervals.

**Figure 4.**
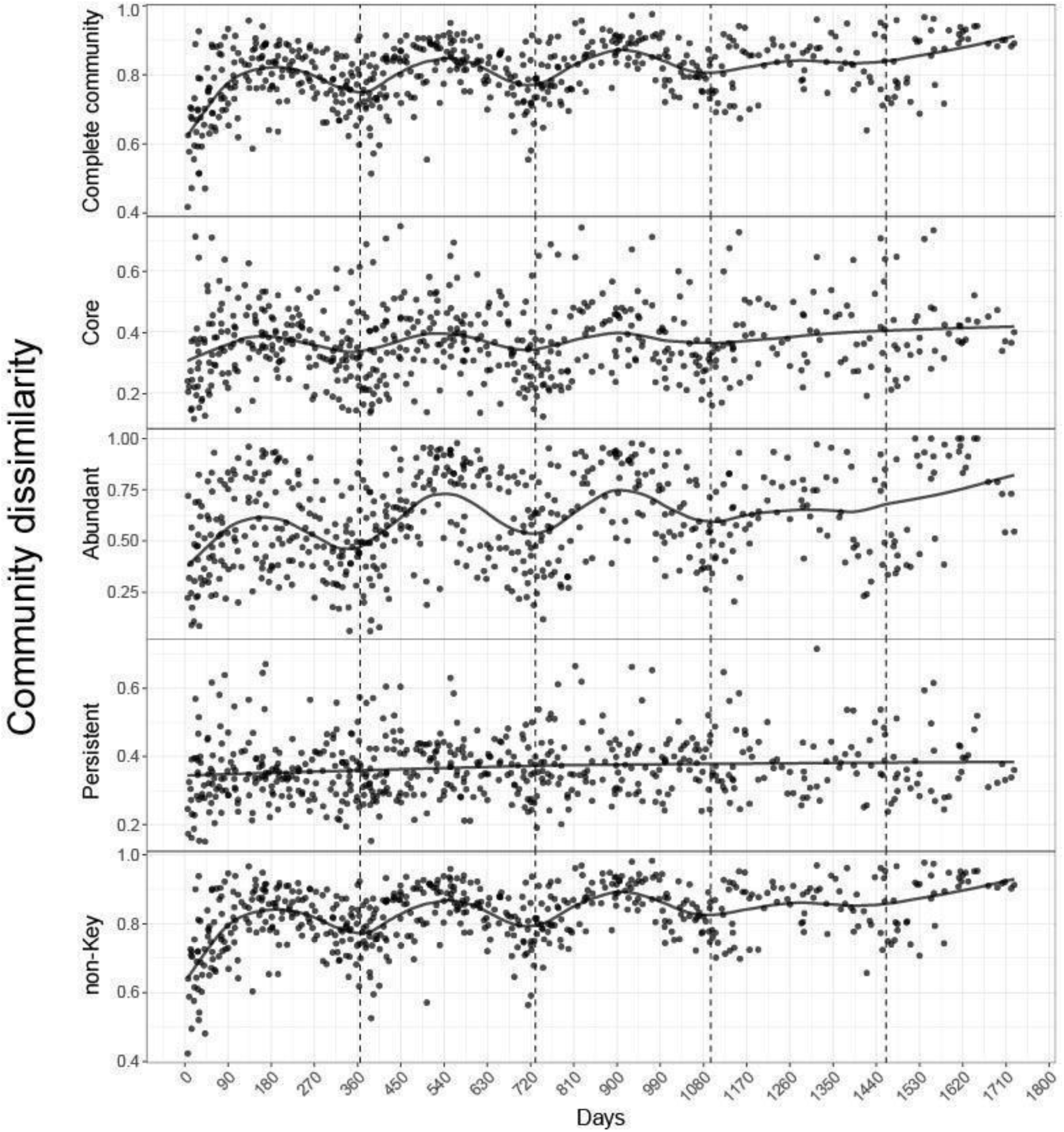
TdR (Time-decay Relationship) plots illustrating temporal variation in sample dissimilarity for the whole community (first box) and, in order, the key groups (Core, Abundant, and Persistent) and non-key groups

The community exhibited higher turnover rates during the transition from Winter-Spring to Summer-Autumn (Fig. 5). This pattern is identical when considering bacteria classified as non-Key, and is similar for the Abundant bacteria, where both gain counts and effective gain values were significantly higher in this direction. In contrast, the pattern was reversed in the Core and Persistent ASV categories For these more stable components of the community, the highest gain values occurred during the transition from summer–autumn to winter–spring (S–A → W–S).

**Figure 5.**
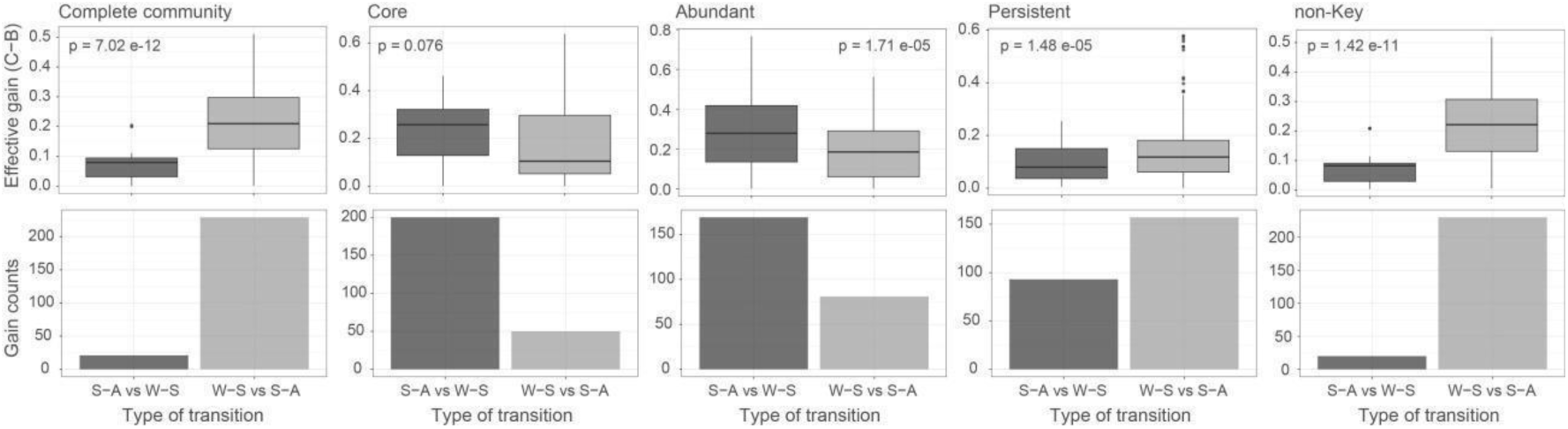
Gains in dissimilarity from the winter-spring transition to summer-autumn, considering, from left to right, the whole community, key groups (Core, Abundant, Persistent), and non-key groups

To explore potential links between picoeukaryotes and prokaryotic dynamics, we analyzed seasonal changes in the relative abundance of 18S-derived ASVs (Fig. S5). The results revealed strong temporal fluctuations in several eukaryotic lineages, particularly Dinoflagellata, and Protalveolata, but these patterns did not align with those observed in the prokaryotic community. Moreover, NMDS analysis of 18S community composition (Fig. S6) indicated a distinct seasonal structuring, with clear separation of spring and summer samples from other seasons. However, this structure was not significantly associated with the measured environmental variables (Fig. S3).

Finally, to understand the paired relationship between the bacteria,eukaryotic ASVS, and CAZymes occurrence, we applied a co-occurrence analysis. The bacterial taxa exhibiting the highest number of interactions with both 16S, 18S, and CAZymes were SAR11, Flavobacteriaceae, and Thiomicrospirales (Table 1; Table S1). SAR11 and Flavobacteriaceae also stood out as the most connected nodes in this co-occurrence network (Fig. 6), which includes additional bacterial representatives such as *Synechococcus, Balneolales, Rodobacterales and Cellvibrionales* among others. The network also reveals a partially isolated substructure formed predominantly by *SAR11* and *Synechococcus*.

**Figure 6.**
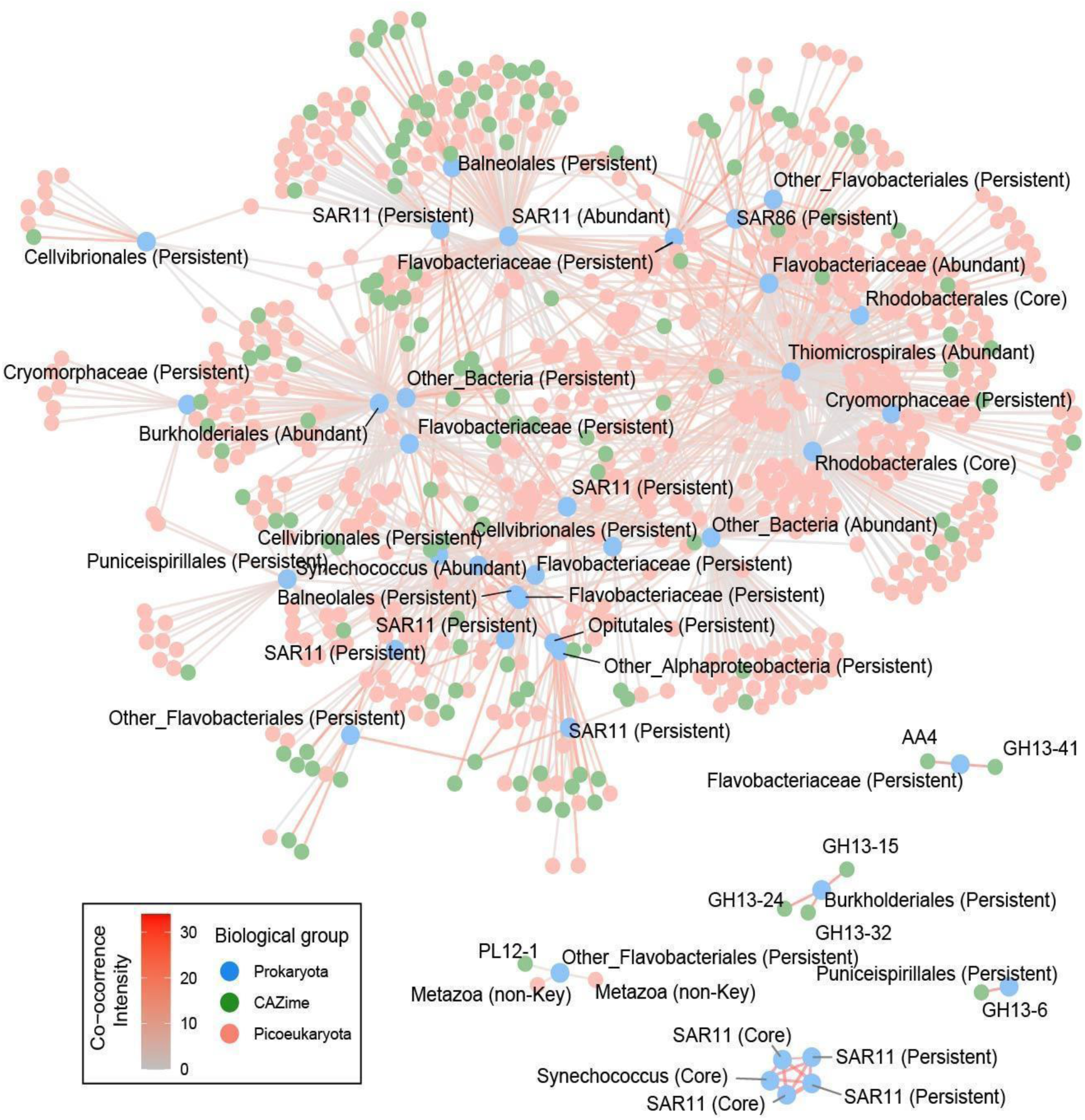
Co-occurrence network illustrating the relationship between bacteria from the key groups (Core, Abundant, and Persistent) among themselves and with CAZymes and picoeukaryotes. For a complete image including the correlations between CAZymes and picoeukaryotes, see Figure S7

**Table 1.**
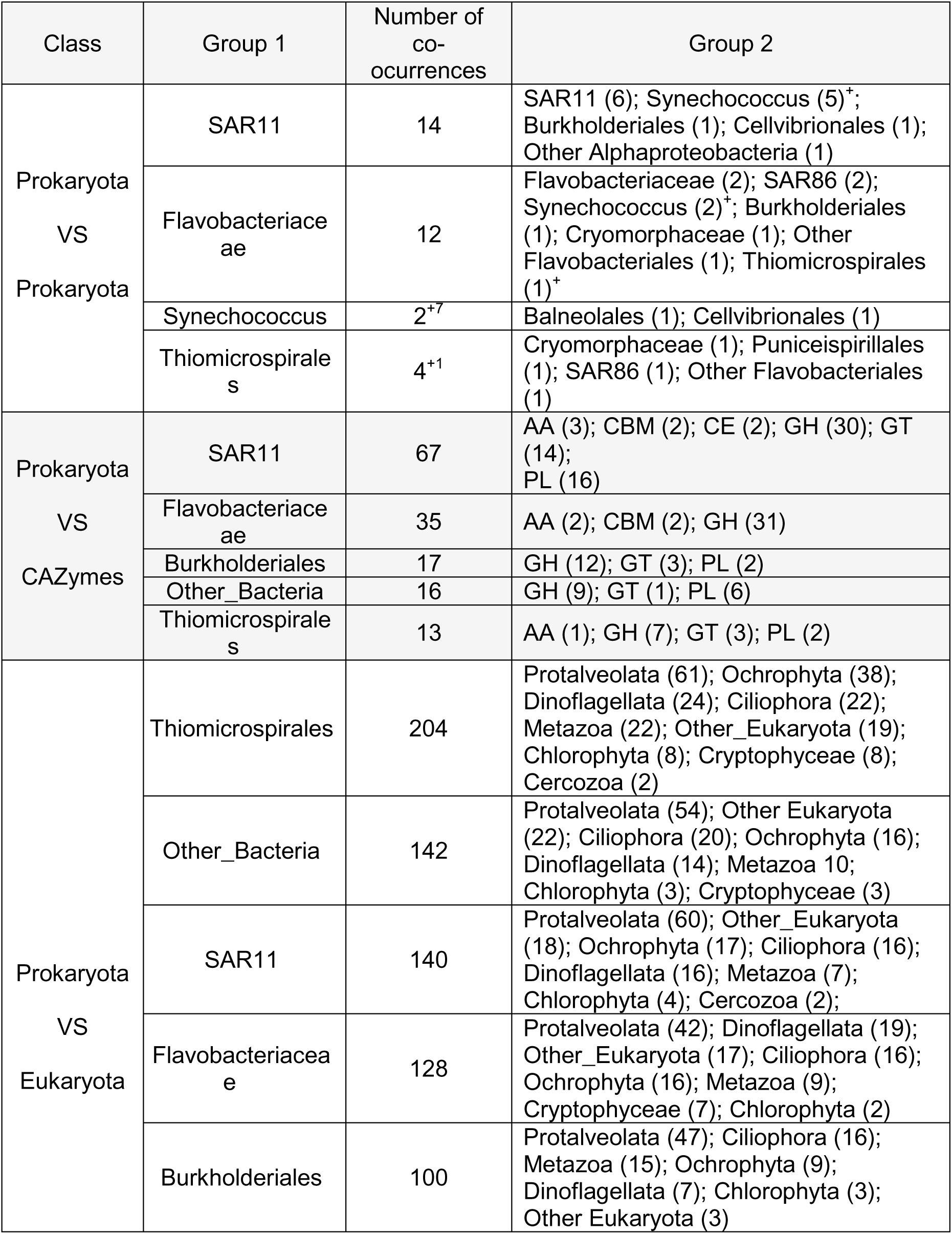
Top 5 bacteria among key groups (Core, Abundant, and Persistent) with the most significant co-occurrences with other bacteria, CAZymes, and picoeukaryotes. For a complete list, see Table S1.

## Discussion

Bacteria are pivotal organisms for understanding ecological processes occurring globally (Chiriac et al. 2022). While their responses to environmental factors have been extensively documented (Lindström et al. 2005, Sunagawa et al. 2015), other drivers such as geographic patterns (Zorz et al. 2019, Mateus-Barros et al. 2021) and biological interactions (Giovannoni 2017, Kim et al. 2019), are gaining increasing recognition, particularly following the advent of culture-independent sequencing technologies (Rappé and Giovannoni 2003). These advancements have unveiled complex and previously unrecognized metabolic dependencies, highlighting theimportance of microbial interactions in shaping community function (Giovannoni 2017). At the same time, the application of classical ecological models provides a robust theoretical framework for interpreting microbial diversity and dynamics (Horner-Devine and Bohannan 2006). Here, we employ a straightforward ecological model — the core-satellite species hypothesis (Hanski 1982) — to identify patterns of persistence over an extended temporal series and to pinpoint groups of potential ecological relevance.

The theory presented by Hanski (1982) posits that, in an ideal sampling, it would be possible to observe a bimodal distribution in the occupancy frequency of any given community, with a large group of organisms being found in all sites. This pattern would be related to the observable abundances for each species within the community, as the most abundant organisms are also those capable of dispersing more efficiently across the landscape (Brown 1984), in a phenomenon called as rescue effect (Gotelli 1991). In this case, in this local study of large-scale temporal span, we sought to understand whether this spatial pattern was also reflected in temporal processes, in a pattern that would reflect the ability of the more resilient bacteria to persist over time and, thus, be those most likely to eventually dominate the (meta)community (Leibold et al. 2004). However, this bimodal pattern could not be observed in the persistence distribution over multiple years. A study conducted in the Baltic Sea (Lindh et al. 2017) demonstrated the existence of a bimodal pattern in the marine bacterioplankton occupation frequency, also related to the high abundance of organisms found at most sites. Although a bimodal pattern in frequency persistency was not directly measured at these sites, the recurrent spatial pattern and repeated finding of common representatives throughout the study, suggest that this could likely be observed. In this case, a relevant geographical feature that may have contributed to it is a more pronounced geographical isolation, which would hinder the action of climate factors on a continental scale while facilitating the action of species sorting (Leibold et al. 2004) by environmental factors, thus guiding these communities.

Here, we have a sampling site highly connected to the Atlantic Ocean, which undergoes successive climate changes and strong selective pressure associated with variations in temperature, salinity, and nutrient input from different sources, which influenced strongly the temporal dynamics of the community. Two distinct groups emerge, clearly and consistently separating temporal variation and grouping characteristic patterns into Winter-Spring and Summer-Autumn. The first is strongly associated with increased local salinity and, more weakly, with a decrease in temperature, and is characterized by a decline in diversity, evenness, and dissimilarity indices. The second appears to be more strongly related to increased local trophic state and, to a lesser extent, to rising temperatures, and is marked by a recurring increase in diversity, evenness, and dissimilarity. These findings are consistent with a previous study conducted at the same site (Pereira et al., 2025). In that study, the authors demonstrated that this pattern is closely related to the seasonal influence of the Brazil and Malvinas currents (i.e. Brazil-Malvinas Confluence; Burone et al. 2021). The Brazil current performing a higher influence during the Summer-Autumn period, is notable for raising the water temperatures and salinity in the region (Burone et al. 2021). This factor favors the increment of biodiversity, but also seems to decrease the functional diversity in this region (Pereira et al., 2025). On the other hand, the colder waters brought by the Malvinas current, particularly during the Winter-Spring period, result in a profound change in local characteristics, which creates challenges for a large portion of the organisms that make up the local bacterioplankton to manage to persist through time, and favors the recurrence of cyanobacteria blooms that occur every spring in the region.

Although a significant bimodality in bacterial frequency persistency cannot be observed, there is still a certain number of ASVs that could be found in all the samples here analyzed. Therefore, we applied an approach that made it possible to identify organisms that may play key roles in local processes. We used the proposed cutoffs to identify the ―Abundant‖ (i.e. relative abundance greater than 1% of the total; Pedrós-Alió 2012) and ―Persistent‖ ASVs (i.e. persistence greater than 90%, similar to the proposed occupancy frequency; Hanski 1982). We also identified a ―Core‖ of bacteria (abundance greater than 1% and persistence greater than 90%) and referred to the remaining community as “non-Key” bacteria. Although each group exhibited its own variation, a recurring pattern across all groups was the association between this variation and local temperature fluctuations and nutrient availability from different sources. This pattern reinforces the link with large-scale changes driven by the Brazil-Malvinas currents confluence, and aligns with other studies in the global ocean indicating that temperature is a predominant factor guiding microbial diversity across the globe (Sunagawa et al. 2015, Zorz et al. 2019).

Among the organisms most commonly found among the key taxa, SAR11 and Flavobacteriaceae stood out. Notably, Flavobacteriaceae are well known for their adaptive capacity, being found in different environments (Chiriac et al. 2022). These organisms possess a complex genome capable of decoding proteins that degrade a wide range of molecules, including CAZymes (Chiriac et al. 2022). Considering the nutrient-limited conditions of the oceanic environment, this ability to degrade carbon compounds of varying complexity may explain their consistent presence over time. In contrast, SAR11 is known for having a reduced genome (i.e., streamlined organisms). A growing effort in the genomic mapping of streamlined organisms has led to significant discoveries regarding these fascinating microbes. Bacteria currently identified only by their genetic codes, such as hgcI in freshwater environments (Kim et al. 2019) and SAR11 in marine environments (Giovannoni 2017), have had their metabolic pathways mapped. This has allowed the identification of molecules essential for their survival but that they cannot synthesize themselves due to the absence a set of essential genes for producing the required proteins. These findings enabled their cultivation in laboratory by the synthesis and provision of these molecules (Giovannoni 2017, Kim et al. 2019). Beyond being a major step toward species-level identification of these groups, this discovery also raises an important ecological question. The presence of these organisms in a given environment implies that these molecules are naturally synthesized—either through the physical breakdown of more complex molecules or via metabolic processing and bioavailability facilitated by other organisms. Moreover, it becomes evident that the availability of these resources may act as a limiting factor for biogeographical processes at both medium and large scales. In the specific case of SAR11, genomic mapping has identified a deficiency in the production of 4-amino-5-hydroxymethyl-2-methylpyrimidine, which can be found in isolates from marine cyanobacterial (Giovannoni 2017). This alone serves as strong evidence of SAR11‘s dependency on other organisms. Additionally, here we identified a recurring fluctuation in their abundance over the years, with an increasing abundance shortly after the annual Synechococcus bloom events and then gradually declining until the following year‘s bloom event. Co-occurrence analyses further support this idea, showing that distinct ASVs identified as belonging to these two groups repeatedly co-occur, even forming a cluster in the co-occurrence network composed exclusively of ASVs from these two taxa.

## Conclusions

By using a simple tool for classifying ASVs among distinct biological components, we had a simpler way to identify groups of interest and factors that guide the variation of diversity as a whole. Here, we were able to demonstrate that the impact of periodical changes in current dynamics seems to be the most relevant factor to explain temporal variations in this community, by influencing the local availability of nutrients and local temperature fluctuations, factors widely recognized as guiding bacterioplankton diversity in the global ocean (Sunagawa et al. 2015). Still, we managed to highlight co-occurrence patterns between some biological components, in a relationship that appears to be mediated by the sharing of organic compounds (Giovannoni 2017). Overall, this approach shows an interesting potential by separating distinct biological components into groups that exhibit ecological coherence, facilitating the identification of processes occurring within this community. Thus, it proves to be an interesting tool for identifying factors guiding their dynamics, both spatially, as previously demonstrated, and temporally.

## Acknowledgements

This project was funded by the ANII-MPI grant (ANII_MPI_ID_2017_1_1007663) awarded to C.A. and R.A., the full-time dedication program (DT) of the Universidad de la República, and PEDECIBA support to C.A. The National Agency for Research and Innovation (ANII) is also acknowledged for supporting E.M.B. as a postdoctoral fellow during the execution of this study (PD_NAC_2023_1_177028). We further acknowledge the support of the Scientific Research Commission of the Universidad de la República Uruguay (CSIC) through the Grupos I+D 2022 program, which contributed to the completion of this work. Special thanks are extended to Danilo Calliari, Maite Colina, Carolina Crisci, Carolina Lescano, Ana Martínez, Mariana Meerhoff, Andrés Pérez, Lorena Rodríguez, Nicolás Silvera, and the personnel from La Paloma Naval Prefecture for their dedication to maintaining the SAMO observatory time series, which is part of the microbial observatories initiative within the µSudAqua network.

