## Supplementary material for "Identification of key bacteria for ecological dynamics in a coastal marine observatory"

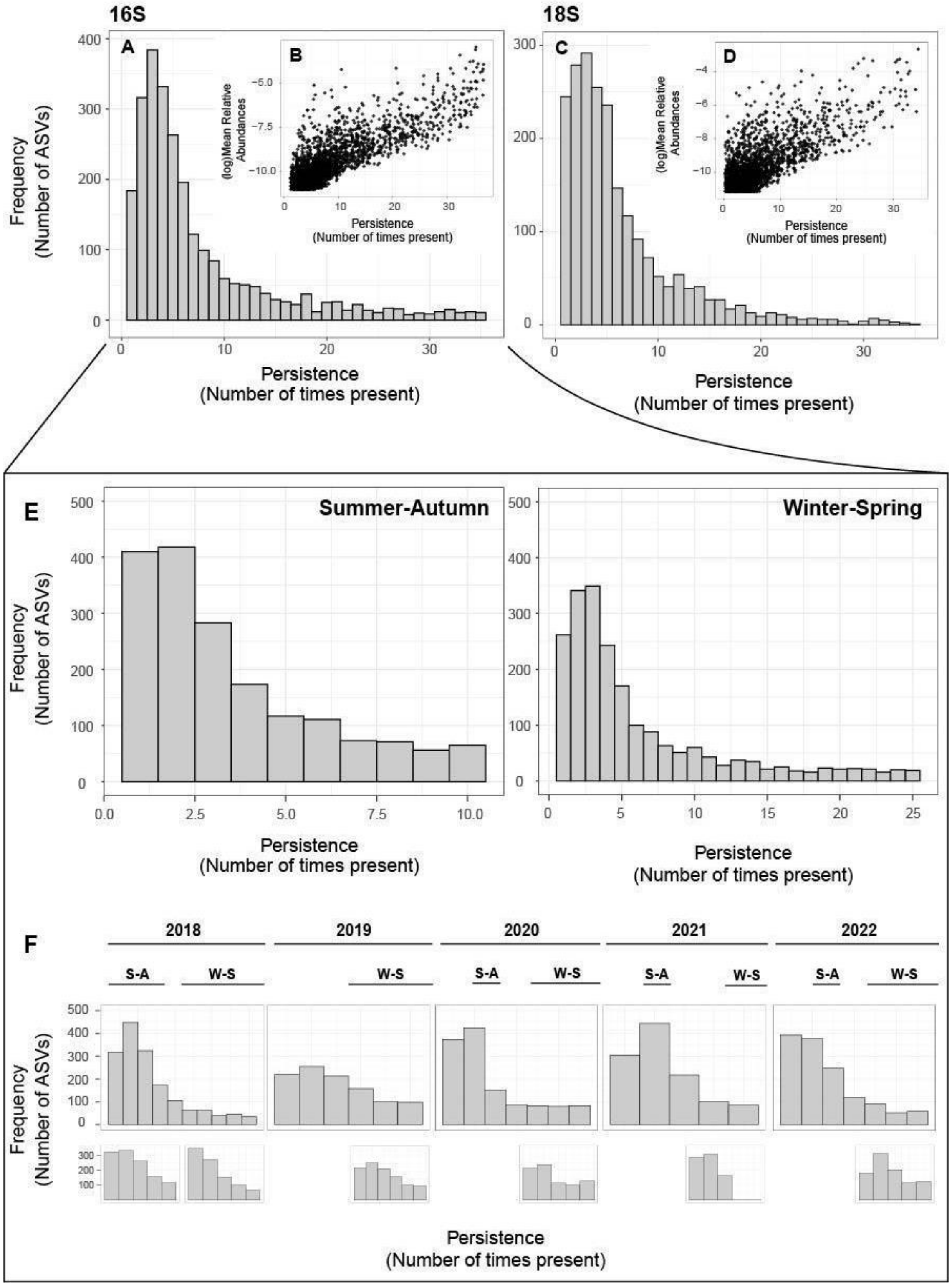


**Figure S1.** Persistence-frequency (A) and mean abundance-frequency (B) distributions of prokaryotic organisms found in 35 samples collected in SAMO between 2018 and 2022. The boxes C and D show the same results, under the same conditions, for eukaryotic organisms. The persistence-frequency distribution analysis of prokaryotic organisms was then broken down to analyze the winter-spring to summer-autumn transition by the Brazil-Malvinas Confluence. Box E shows the results considering all samples from all years collected during the winter-spring or summer-autumn period, while box F shows the results when considering samples collected during the winter-spring or summer-autumn period for each year. The Mitchell-Olds & Shaw test showed a significant result ($p<0.05$). The upper boxes show values for prokaryotes, while the bottom boxes show the same results for eukaryotes

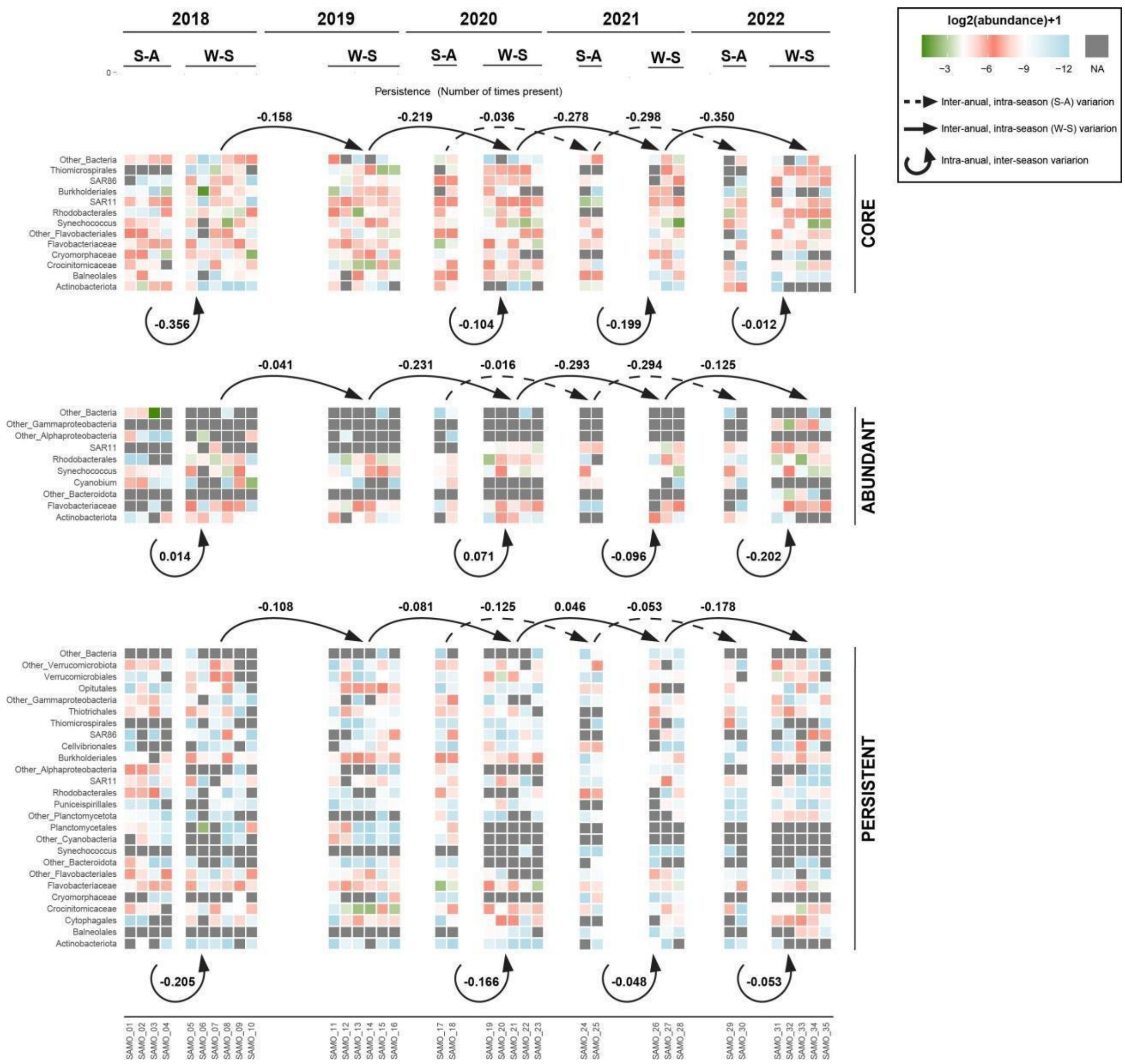


**Figure S2.** Heatmap showing changes in the abundance of the groups of interest across the winter-spring or summer-autumn periods for each year, considering the Core, Abundant, and Persistent bacteria from top to bottom. Arrows show the magnitude of the observed variation between periods

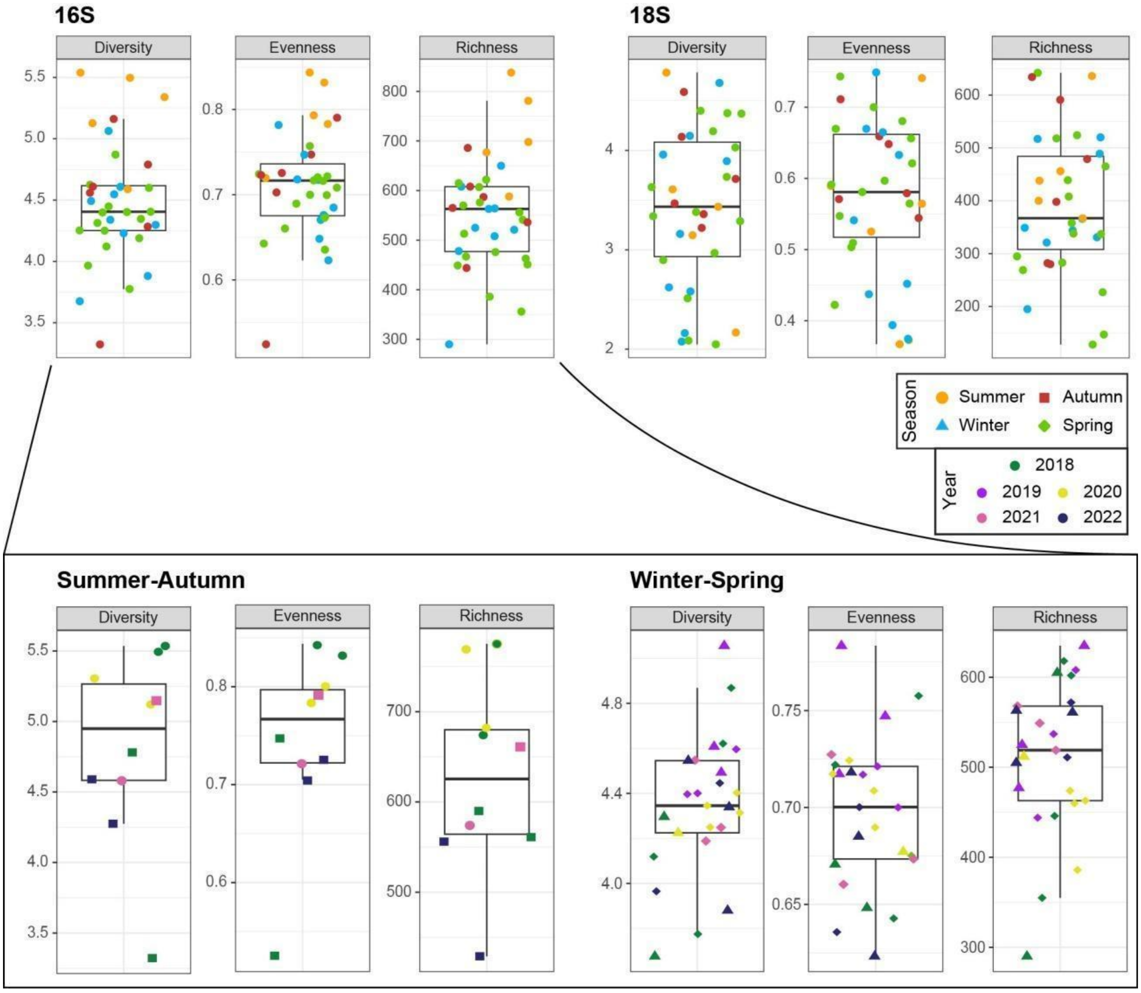


**Figure S3.** Box plots displaying values of diversity, evenness, and richness for prokaryotes (upper boxes on the left) and eukaryotes (upper boxes on the left). Each point was colored to indicate the season in which the respective sample was collected. In the lower box, the samples were grouped by season, considering the winter-spring to summer-autumn transition by the Brazil-Malvinas Confluence. Here, the shapes represent the season in which each sample was collected, and the colors represent the year of collection

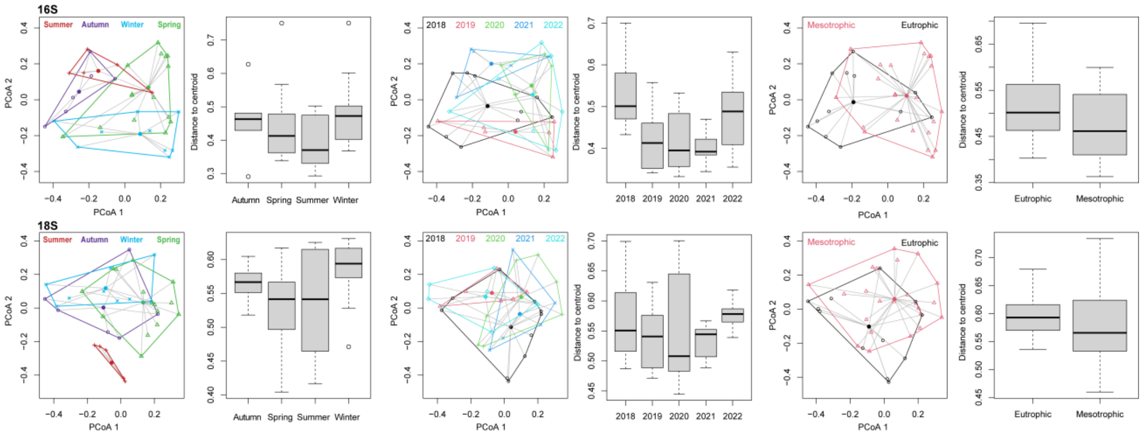


**Figure S4.** Beta-dispersion plots showing the intra-group dissimilarity considering, from left to right, the seasons, the year of collection, the winter-spring to summer-autumn transition by Brazil-Malvinas Confluence, and trophic state for prokaryotes (upper boxes) and eukaryotes (lower boxes)

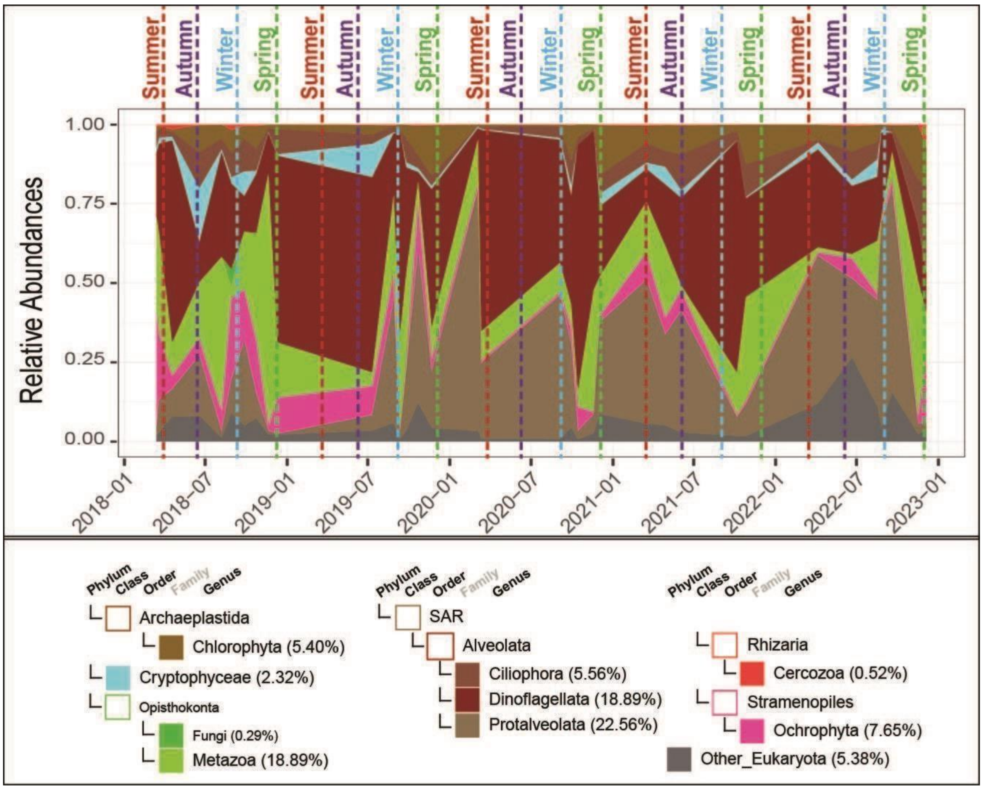


**Figure S5.** Temporal variation in the relative abundances of phytoplanktonic picoeukaryotes. Each ASV was classified according to its phylogenetic phylum. Groups of known relevance for local dynamics were classified at the Class or Order level, depending on the classification refinement for each phylum. The values in parentheses indicate the total relative abundance of each group

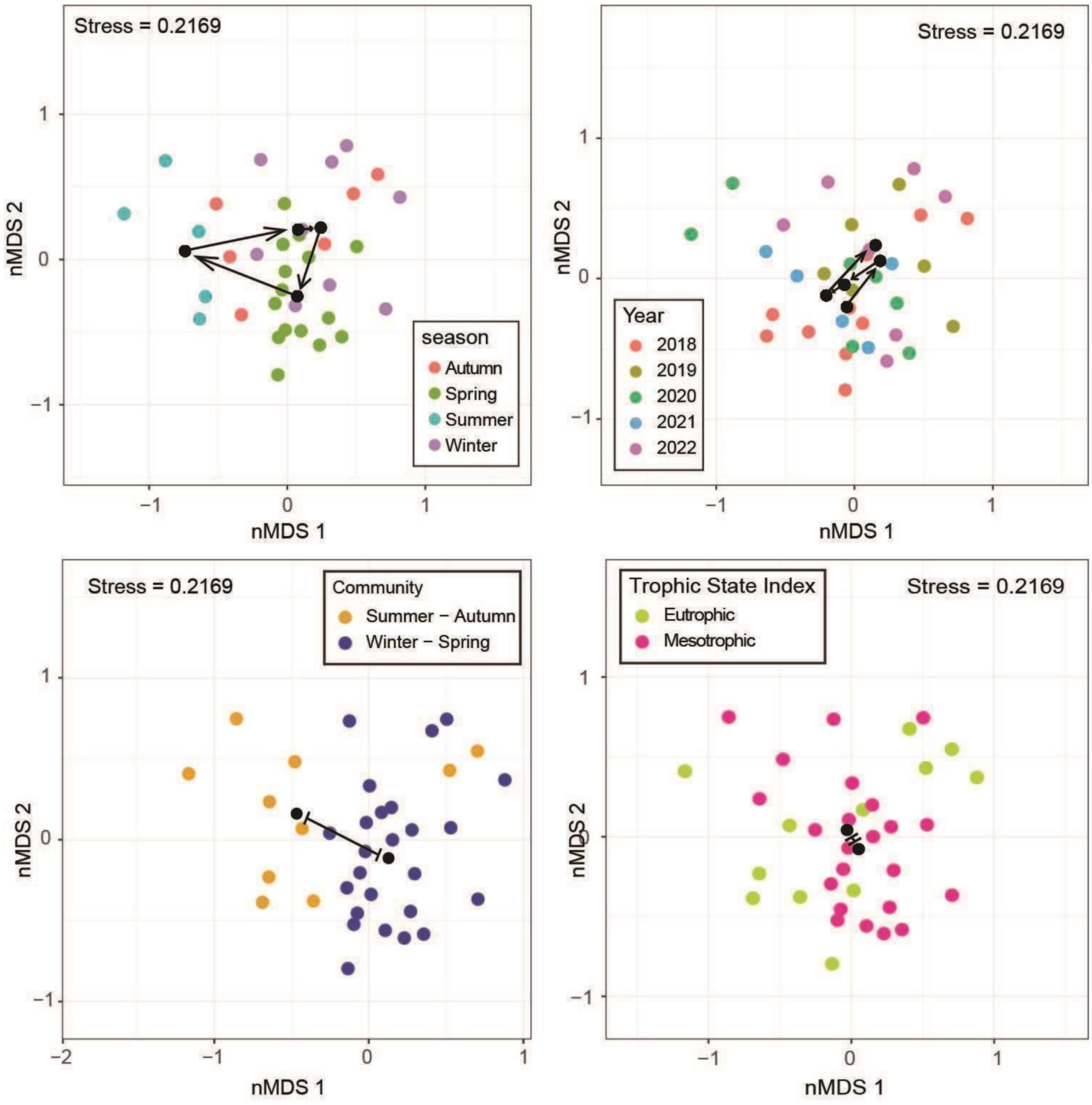


**Figure S6.** nMDS (non-metric multidimensional scaling) plots showing the dissimilarity in planktonic picoeukaryotes considering large-scale environmental changes across seasons (upper left box), year of collection (upper right box), the winter-spring to summer-autumn transition at the Brazil-Malvinas Confluence (lower left box), and trophic state (lower right box)

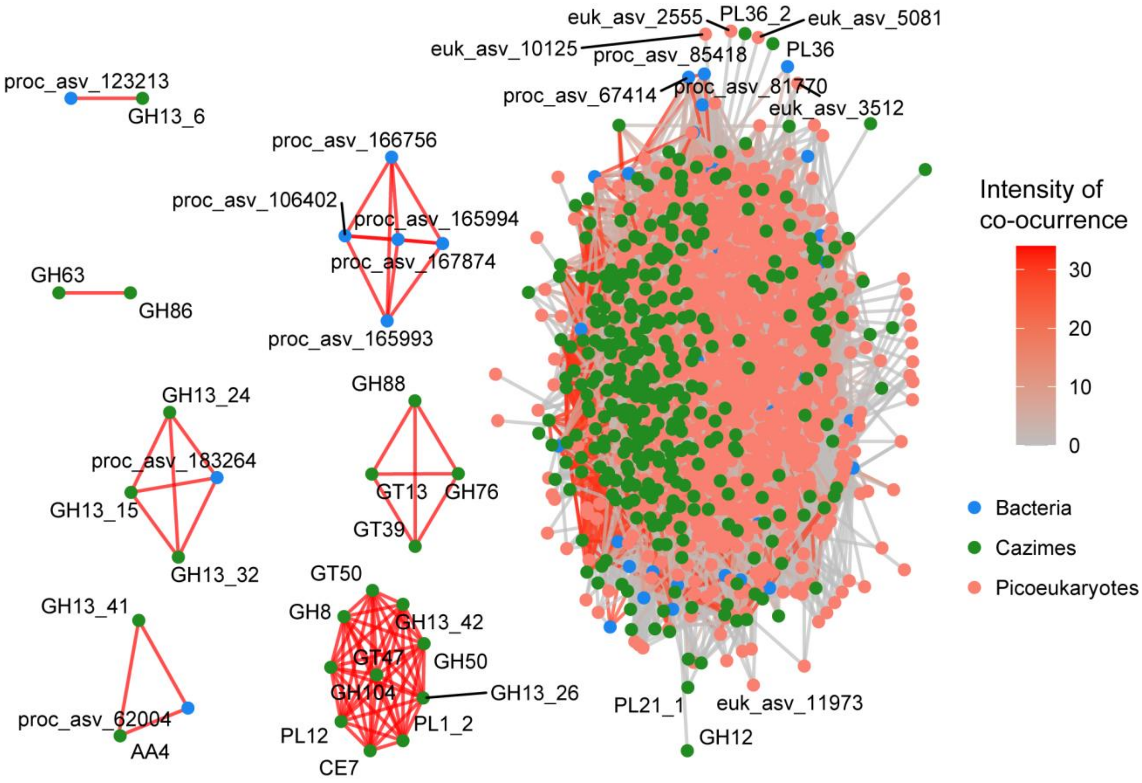


**Figure S7.** Co-occurrence network displaying the relationship between the bacteria from the key group (Core, Abundant, and Persistent), CAZymes, and picoeukaryotes

**Table S1.** Bacteria from the key groups (Core, Abundant, and Persistent) that exhibited the most significant co-occurrences with other bacteria, CAZymes, and picoeukaryotes.

| Class | Group 1 | Number of co-ocurrences | Group 2 |
|---|---|---|---|
| Prokaryota VS Prokaryota | SAR11 | 14 | SAR11 (6); Synechococcus (5)$^{+}$; Burkholderiales (1); Cellvibrionales (1); Other Alphaproteobacteria (1) |
| | Flavobacteriaceae | 12 | Flavobacteriaceae (2); SAR86 (2); Synechococcus (2)$^{+}$; Burkholderiales (1); Cryomorphaceae (1); Other Flavobacteriales (1); Thiomicrospirales (1)$^{+}$ |
| | Synechococcus | $2^{+7}$ | Balneolales (1); Cellvibrionales (1) |
| | Thiomicrospirales | $4^{+1}$ | Cryomorphaceae (1); Puniceispirillales (1); SAR86 (1); Other Flavobacteriales (1) |
| Prokaryota VS CAZimes | SAR11 | 67 | AA (3); CBM (2); CE (2); GH (30); GT (14); PL (16) |
| | Flavobacteriaceae | 35 | AA (2); CBM (2); GH (31) |
| | Burkholderiales | 17 | GH (12); GT (3); PL (2) |
| | Other_Bacteria | 16 | GH (9); GT (1); PL (6) |
| | Thiomicrospirales | 13 | AA (1); GH (7); GT (3); PL (2) |
| | Cellvibrionales | 11 | CE (1); GH (8); GT (1); PL (1) |
| | Balneolales | 11 | AA (1); GH (7); GT (2); PL (1) |
| | Opitutales | 9 | AA (1); CBM (1); GH (2); GT (1); PL (4) |
| | Other Alphaproteobacteria | 8 | AA (1); GH (4); GT (3) |
| | Other Flavobacteriales | 8 | GH (3); GT (2); PL (3) |
| | Rhodobacterales | 7 | CBM (2); CE (1); GH (1); PL (3) |
| | Synechococcus | 7 | AA (1); GH (5); GT (1) |
| | Thiomicrospirales | 204 | Protalveolata (61); Ochrophyta (38); Dinoflagellata (24); Ciliophora (22); Metazoa |

| | | | |
|---|---|---|---|
| Prokaryota VS Eukaryota | | | (22); Other_Eukaryota (19); Chlorophyta (8); Cryptophyceae (8); Cercozoa (2) |
| | Other_Bacteria | 142 | Protalveolata (54); Other Eukaryota (22); Ciliophora (20); Ochrophyta (16); Dinoflagellata (14); Metazoa 10; Chlorophyta (3); Cryptophyceae (3) |
| | SAR11 | 140 | Protalveolata (60); Other_Eukaryota (18); Ochrophyta (17); Ciliophora (16); Dinoflagellata (16); Metazoa (7); Chlorophyta (4); Cercozoa (2); |
| | Flavobacteriaceae | 128 | Protalveolata (42); Dinoflagellata (19); Other_Eukaryota (17); Ciliophora (16); Ochrophyta (16); Metazoa (9); Cryptophyceae (7); Chlorophyta (2) |
| | Rhodobacterales | 100 | Protalveolata (47); Ciliophora (16); Metazoa (15); Ochrophyta (9); Dinoflagellata (7); Chlorophyta (3); Other Eukaryota (3) |
| | Puniceispirillales | 73 | Protalveolata (29); Ochrophyta (12); Dinoflagellata (9); Other Eukaryota (7); Ciliophora (6); Cryptophyceae (4); Metazoa (4); Chlorophyta (2) |
| | Burkholderiales | 60 | Protalveolata (25); Other Eukaryota (19); Dinoflagellata (5); Metazoa (5); Chlorophyta (3); Ciliophora (3) |
| | Synechococcus | 58 | Protalveolata (28); Ochrophyta (10); Other Eukaryota (9); Ciliophora (8); Dinoflagellata (3) |
| | Cellvibrionales | 46 | Protalveolata (18); Ochrophyta (8); Ciliophora (7); Other Eukaryota (6); Metazoa (4); Dinoflagellata (3) |
| | Cryomorphaceae | 32 | Ochrophyta (11); Ciliophora (5); Cryptophyceae (5); Chlorophyta (4); Other_Eukaryota (4); Dinoflagellata (3) |
| | Other Flavobacteriales | 18 | Ciliophora (6); Ochrophyta (4); Protalveolata (4); Cryptophyceae (2); Metazoa (2) |
| | | 14 | Other_Eukaryota (5); Dinoflagellata (4); Protalveolata (3); Metazoa (2) |
| | Balneolales | 10 | Protalveolata (8); Ciliophora (2) |
| | Other Alphaproteobacteria | 10 | Ciliophora (3); Dinoflagellata (3); Ochrophyta (2); Other_Eukaryota (2) |
| | Opitutales | 2 | Other_Eukaryota (2) |
